# The crystal structure of nsp10-nsp16 heterodimer from SARS-CoV-2 in complex with S-adenosylmethionine

**DOI:** 10.1101/2020.04.17.047498

**Authors:** Monica Rosas-Lemus, George Minasov, Ludmilla Shuvalova, Nicole L. Inniss, Olga Kiryukhina, Grant Wiersum, Youngchang Kim, Robert Jedrzejczak, Natalia I. Maltseva, Michael Endres, Lukasz Jaroszewski, Adam Godzik, Andrzej Joachimiak, Karla J. F. Satchell

## Abstract

SARS-CoV-2 is a member of the coronaviridae family and is the etiological agent of the respiratory Coronavirus Disease 2019. The virus has spread rapidly around the world resulting in over two million cases and nearly 150,000 deaths as of April 17, 2020. Since no treatments or vaccines are available to treat COVID-19 and SARS-CoV-2, respiratory complications derived from the infections have overwhelmed healthcare systems around the world. This virus is related to SARS-CoV-1, the virus that caused the 2002-2004 outbreak of Severe Acute Respiratory Syndrome. In January 2020, the Center for Structural Genomics of Infectious Diseases implemented a structural genomics pipeline to solve the structures of proteins essential for coronavirus replication-transcription. Here we show the first structure of the SARS-CoV-2 nsp10-nsp16 2’-O-methyltransferase complex with S-adenosylmethionine at a resolution of 1.80 Å. This heterodimer complex is essential for capping viral mRNA transcripts for efficient translation and to evade immune surveillance.

## Introduction

On December 31^st^, 2019, the World Health Organization (WHO) was alerted of a pneumonia outbreak with an unknown etiology, originating in the Chinese province of Wuhan, Hubei (1,2). The etiological agent was identified as a coronavirus, closely related to the virus responsible for Severe Acute Respiratory Syndrome (SARS). The new SARS coronavirus-2 (SARS-CoV-2) causes the severe respiratory infection, Coronavirus Disease 2019 (COVID-19) (3). Within four months, SARS-CoV-2 rapidly spread, sparking a global pandemic. The COVID-19 pandemic has also forced government-enacted “stay-at-home” orders around the world. According to the World Health Organization, 2,074,529 SARS-CoV-2 infections have been confirmed, of which 139,378 were fatal as of April 17^th^. These data are similar to Johns Hopkins University tracking system, that reported 2,182,734 of infections and 147,384 deaths (1).

The coronaviridae family of viruses causes disease in birds and mammals, including bats, camels, pigs, and humans. In lower vertebrates, pathogenic coronaviruses cause acute and severe gastrointestinal infections, fevers and organ failure. Of the seven human-tropic coronaviruses, hCoV-229E, hCoV-NL63, hCoVB-OC43 cause only asymptomatic or mild infections, including the common cold (4-6). Four of the viruses are linked to severe infections; including, hCoV-HKU1, a common cause of pneumonia, SARS-CoV-1 with a 10% mortality rate, Middle East Respiratory Syndrome Virus (MERS-CoV) with a 37% mortality rate (3), and SARS-CoV-2 currently with a 6% mortality rate for confirmed cases (1). As SARS-CoV-2 continues to spread, the need for effective vaccines and therapeutics increases. In addition, there is currently little information about the immunological response to the virus or the potential for re-infection (7-10). Therefore, it is urgent to study SARS-CoV-2 mechanisms of infection and replication in order to find effective targets for drug and vaccine development.

Coronaviruses have a large (∼ 30 kb) single-stranded, positive RNA genome that is 5’-capped, contains a 3’-poly-A tail, and are direct templates for the transcription of sub-genomic mRNAs for the translation of viral proteins. The first open reading frame produces the large non-structural polyprotein 1a (pp1a) and read-through across a frameshift results in translation of the larger non-structural polyprotein 1ab (pp1a/b). These polyproteins are subsequently processed into sixteen non-structural proteins (nsps) that assemble to form the Replication-Transcription Complex (RTC) or function as accessory proteins necessary for viral replication. The structural and additional accessory proteins are encoded at 3’-end of the genome (11-14).

The components of the RTC include enzymes that regulate mRNA and genomic RNA synthesis, proofreading, and mRNA maturation. Two of these enzymes are critical for capping viral mRNAs, a tactic employed by multiple RNA viruses to avoid immune detection by toll-like receptors 7 (TLR7) and 8 (TLR8) (15). In eukaryotic cells, mRNA capping is initiated by an RNA triphosphatase (TPase), which removes the γ-phosphate from the 5’-end of the nascent mRNA transcript, generating a diphosphate 5’-ppN end. An RNA guanylyltransferase (GTase) subsequently catalyzes the hydrolysis of pyrophosphate (PPi) from a guanidine triphosphate (GTP) molecule forming GMP, followed by the transfer of the α-phosphate of guanidine monophosphate (GMP) to the diphosphate 5’-ppN transcript end, forming the cap core structure, referred to as GpppN. The GpppN formation is followed by N^7^-methylation of the capping guanylate by a guanine-N^7^-methyltransferase (N^7^-MTase) to generate the Cap-0. Further methylation at the ribose 2’-O position of first nucleotide of the RNA is catalyzed by a ribose 2’-O-methyltransferases (2’-O-MTase) to generate Cap-1 and sometimes at the second nucleotide to generate Cap-2 (4). Both the N^7^-MTase and 2’-O-MTase use S-adenosyl-L-methionine (SAM) as the methyl group donor (4,16).

For coronavirus mRNA maturation, the host cell TPases and GTase are used to guanylate the 5’-end of the nascent mRNA and the viral non-structural protein 14 (nsp14) N^7^-MTase activity generates the Cap-0 (4). Nsp14 is a bifunctional enzyme with an exonuclease domain in addition to its N^7^-MTase domain (17). Its activity is modulated by the binding of the small viral protein, nsp10, which specifically stimulates its exonuclease activity with no effect on its N^7^-MTase activity (16). The coronavirus mRNAs are further modified to have a Cap-1 by the viral non-structural protein 16 (nsp16). Nsp16 is a m^7^GpppA-specific, SAM-dependent, 2’-O-MTase (18,19) and is activated by binding to nsp10 (20). Nsp10 is a stable protein that can self-dimerize (5) or form dodecamers when the nsp11 extension is included (6), in addition to modulating nsp14 and nsp16 activity (21). Although no specific enzymatic activity has been identified for nsp10 and its fold is unique, it is known as a zinc binding protein that binds nonspecifically to RNA and stabilizes the SAM binding pocket in nsp16 to form a stable complex (20). The nsp10-nsp16-mediated 2’-O-methylation of coronavirus RNA is essential for preventing host recognition and decreasing immune response, while the viral RNA is translated (16,18,22).

Multiple crystal structures of the nsp16-nsp10 heterodimer have been solved, from both SARS-CoV-1 and MERS-CoV (4,19,23) and provide some insight into the structural basis of substrate binding and the proposed sn2-mechanism of methyl transfer. To facilitate the development of vaccine and drug targets for SARS-CoV-2, the Center for Structural Genomics for Infectious Diseases (CSGID) has implemented a high-throughput structural determination pipeline in order to solve the structure of SARS-CoV-2 proteins that are amenable to drug discovery and vaccine development. Here we present the 1.80 Å X-ray crystal structure of SARS-CoV-2 nsp10-nsp16 heterodimer with SAM bound.

## Results

### Overall structure of SARS-CoV-2 nsp10-nsp16 heterodimer

Nsp10 and nsp16 are encoded in the polycistronic *orf1ab* of the (+)ssRNA from SARS-CoV-2 (Fig. 1A) and are released by processing of the polyproteins by the nsp5 protease (14). Nsp10 is released from pp1a and pp1ab, while nsp16 is part of only the pp1ab polyprotein created after a −1 ribosome shift (24). Previous studies from SARS-CoV-1 demonstrated that nsp16 is an unstable protein in several conditions and thus the protein was purified in complex following co-expression with nsp10 (25). Here, we successfully expressed and purified nsp10 and nsp16 separately using nickel affinity and size exclusion chromatography (SEC). After removal of the 6xHis-tags, a heterodimer of nsp16 with nsp10 was formed and concentrated by centrifugation across a 30 kDa cutoff membrane and dialyzed into crystallization buffer. The protein complex was further incubated with 2 mM SAM and diffraction quality crystals grew in A7 condition of the ComPAS (Qiagen) screen (0.2 M Calcium acetate, 0.1 M HEPES, pH 7.5, 18% w/v PEG 8000).

**Figure 1.**
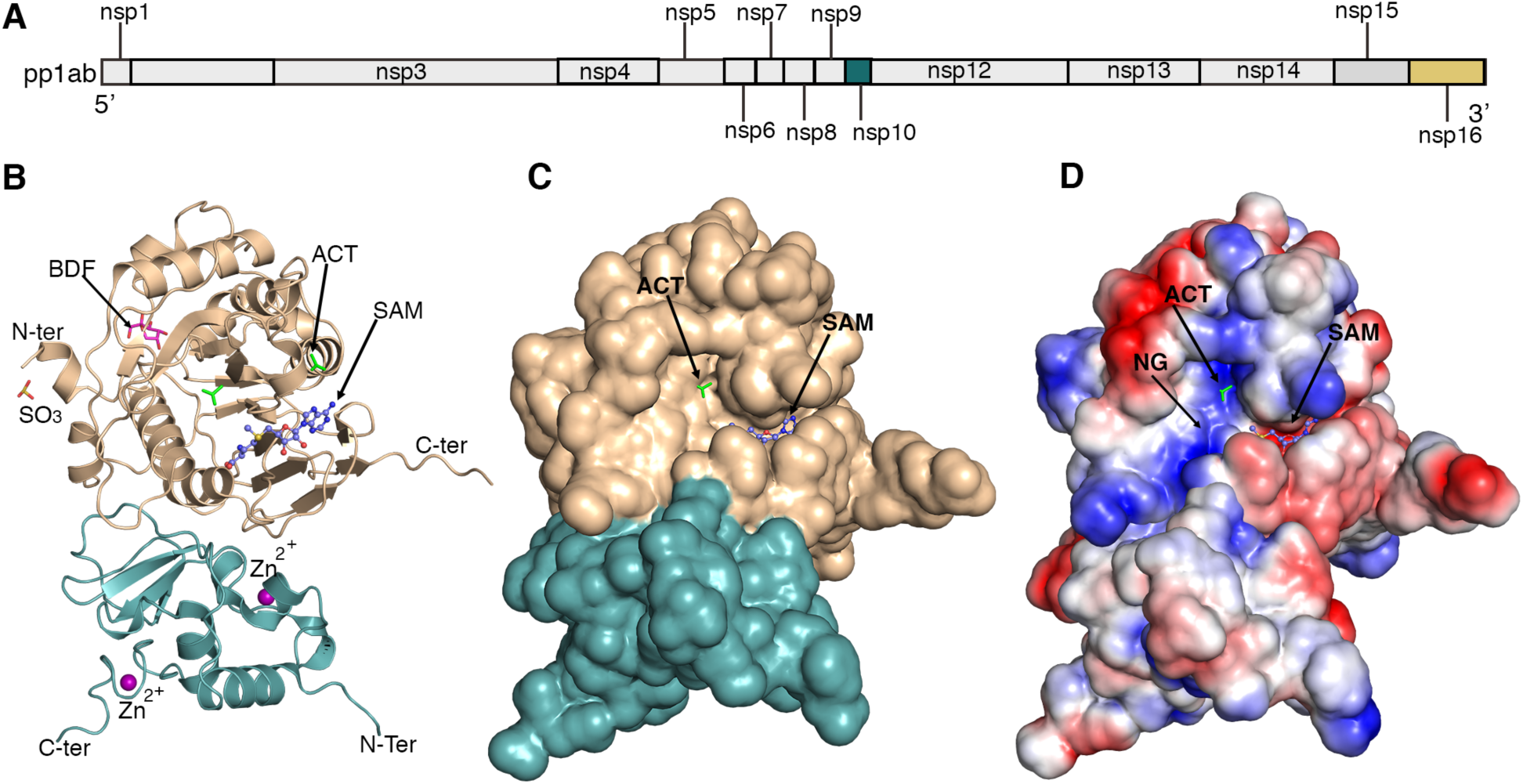
Overall structure of the nsp10-nsp16. A, Linear schematic of the orf1ab translation produced prior to nsp3 and nsp5 processing. B, Cartoon representation of the heterodimer of nsp16 (tan) in complex with nsp10 (teal). Ligands are sulfite (SO_3_), β-D-fructopyranose, (BDF), acetate, (ACT), S-adenosyl-methionine (SAM), and Zn ^2+^. C, Solvent exposed surface, showing the SAM pocket and ACT. D, Surface charge showing the positively charged surface at the groove for nucleotide (NG) binding site and the negatively charged SAM binding pocket (C).

The structure of nsp16 in complex with nsp10 and SAM was solved by molecular replacement at 1.8 Å (PDB code 6w4h) using the nsp10-nsp16 heterodimer structure from SARS-CoV-1 (PDB code 3r24) as the model. The diffraction data are presented in Table 1 and a representative image of the structures is shown in Figure 1. The crystal belonged to the space group P3_1_21 with two polypeptide chains, one chain of nsp10 and one chain of nsp16 in the asymmetric unit, which form a heterodimer. The heterodimer has a total solvent-exposed surface area of 19,710 Å^2^ estimated by PISA (Fig. 1B). This heterodimer is mainly stabilized by hydrophobic interactions and hydrogen bounds in the interface of nsp10 and nsp16. In parallel with this work, a nearly identical structure with the same space group was solved at 2.0 Å (PDB code 6w61). The crystal obtained for this structure was prepared by a distinct procedure (see materials and methods). A third structure was also solved at 1.95 Å (PDB code 6w75) by the same methods used for the first structure. The third crystal form belongs to the P3_2_21 space group with twice the unit cell than first two. In this structure, there are four polypeptide chains, which form a tetramer, and this structure will be described later. For this report, only highest resolution structure (PDB code 6w4h) is discussed in detail.

In the representative crystal structure (PDB code 6w4h), there are several ligands bound to the heterodimer: two molecules of acetate (ACT), two molecules of β-D-fructopyranose (BDF), one molecule of sulfite (SO_3_), two Zinc ions and one molecule of SAM. All of these ligands, except the MTase co-factor SAM and two Zinc ions, are derived from the crystallization and freezing solutions. It is likely that their interaction with the protein complex results from its electrostatic potential, as predicted by the Adaptive Poisson-Boltzmann Solver program (26) and modeled as vacuum surface electrostatics in the software package PyMOL. For example, one of the ACT molecules interacts in the positively charged nucleotide binding groove (Fig. 1C).

### Structure of nsp10

The structure of SARS-CoV-2 nsp10 consist of residues 19-133 (residues 4272-4392 of pp1a). It contains a positively charged and hydrophobic surface that interacts with a hydrophobic pocket and a negatively charged surface from nsp16, which contributes to the stabilization of the SAM binding site (Fig. 1C). The structure of nsp10 of SARS-CoV-2 is composed of a central anti-parallel pair of β-strands surrounded on one side by a crossover large loop. The other side is a helical domain with loops that form two zinc fingers (Figure 2 A). These structures are involved in the non-specific binding of RNA in other coronaviruses (4,27).The Zn-binding site 1 is coordinated by the residues C74, C77, H83 and C90. The Zn-binding site 2 is coordinated by C117, C120, C128, and C130.

**Figure 2.**
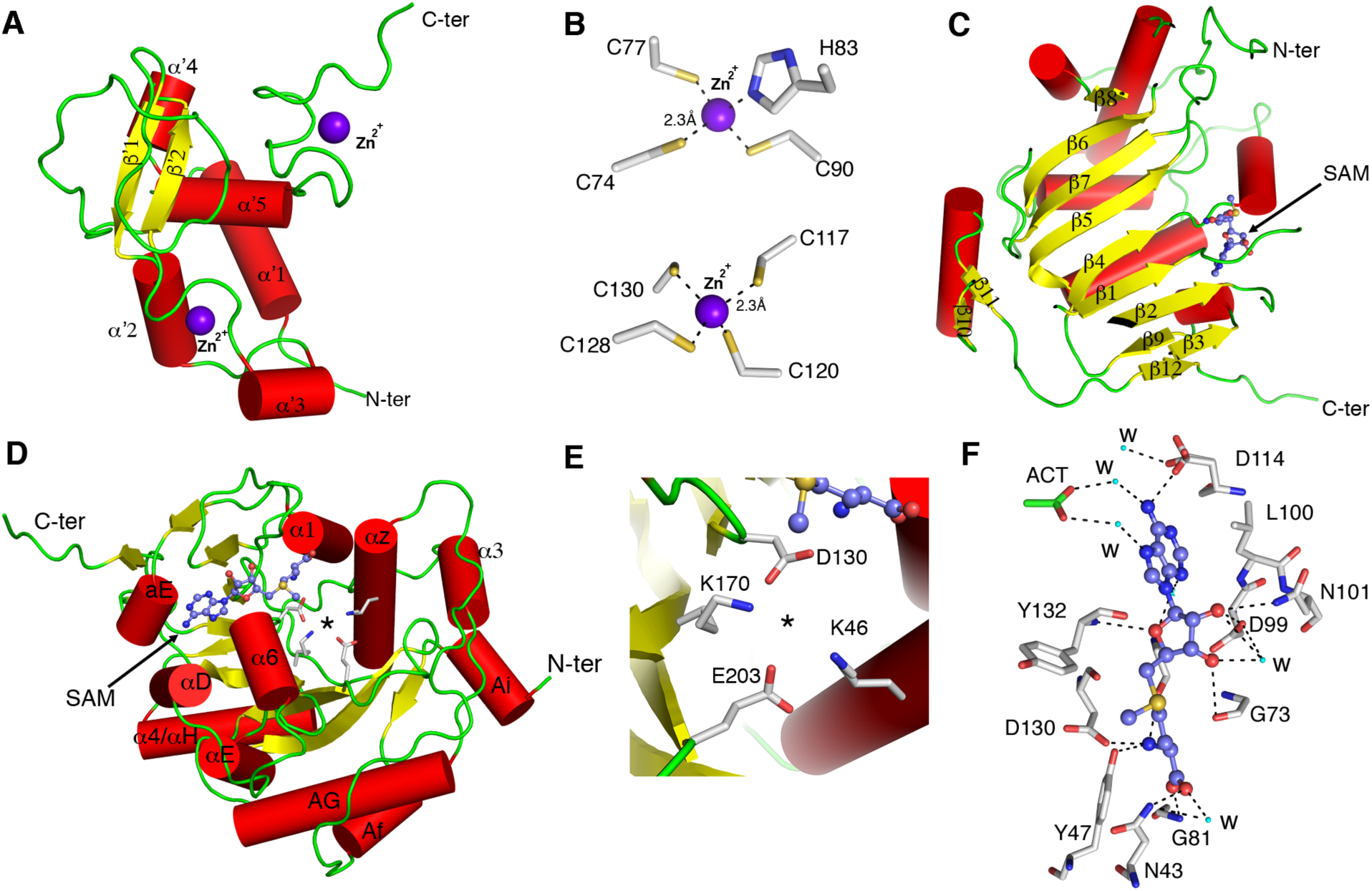
Detailed representation of nsp10 and nsp16 structures and ligands. A, representation of the structure of nsp10 with 5 helices (red) and 2 beta-strands (yellow) and two zinc ions (purple). B, Close-up view of the two Zn^2+^ binding sites in nsp10. C, canonical beta sheet from nsp16 bound with SAM (ball and stick). D, nsp16 helices, SAM (ball and stick) and catalytic residues (gray sticks). Asterisk indicates the catalytic methyl transferase residues. E, Close-up view of the catalytic residues in nsp16. F, Detailed view of coordination of SAM binding. Small cyan dots indicate water molecules (w) and dashed lines the hydrogen bonds. Acetate (ACT) is in green sticks.

### Structure of nsp16

The nsp16 structure has all 298 residues (orf1ab residues 6799-7096) with the Serine-Asparagine-Alanine residues at the N-terminus derived from the tobacco etch virus (TEV) protease cleavage site. The structure of nsp16 contains the 2’-O-MTase catalytic core comprised of a Rossmann-like β-sheet fold decorated by eleven α-helices, seven β-strands, and loops (Fig. 2C and D). The SARS-CoV-2 nsp16 fold is formed by a β-sheet with the canonical 3-2-1-4-5-7-6 arrangement, in which β7 is the only antiparallel strand (Figure 2C). This β-sheet is sandwiched by loops and α-helices. The catalytic core binds one molecule of SAM near the β1 and β2 strands of the Rossmann-like fold, and the negatively charged binding SAM binding pocket is formed by the loops 71-79 and 100-108, and 130-148 (Figure 2D). Residues Lys-46, Asp-130, Lys-170, and Glu-203, comprising the canonical SAM MTase motif, are in close proximity to the SAM methyl group that is catalytically transferred to the 2’-O sugar (Fig. 2 E) (4,16,19,28).

### nsp10 and nsp16 are highly conserved among lineage B of the beta coronaviruses

In order to explore whether SARS-CoV-2 has amino acid variation in the 2-O’MTase that could impact its function, we compared the primary amino acid sequences of nsp10 and nsp16 proteins to that of other beta-coronaviruses (Fig. 3 A and B). Nsp10 was found to be 100% identical to Bat-CoV-RaTG13, 99% identical to SARS-CoV-1, and 98% identical to Bat-SARS-like coronavirus (Bat-SL-CoV) isolate Rs4247, but only 59% identical to MERS-CoV (Figure 3A). In the aligned sequences, the zinc coordinating residues C74, C77, H83, C90, C117, C120, C128, and C130 are 100% conserved, emphasizing the importance of zinc binding for nsp10 activity during the viral replication-transcription processes. Thus, nsp10 is highly conserved in the lineage B beta-coronaviruses, and less conserved with lineage C beta coronaviruses.

**Figure 3.**
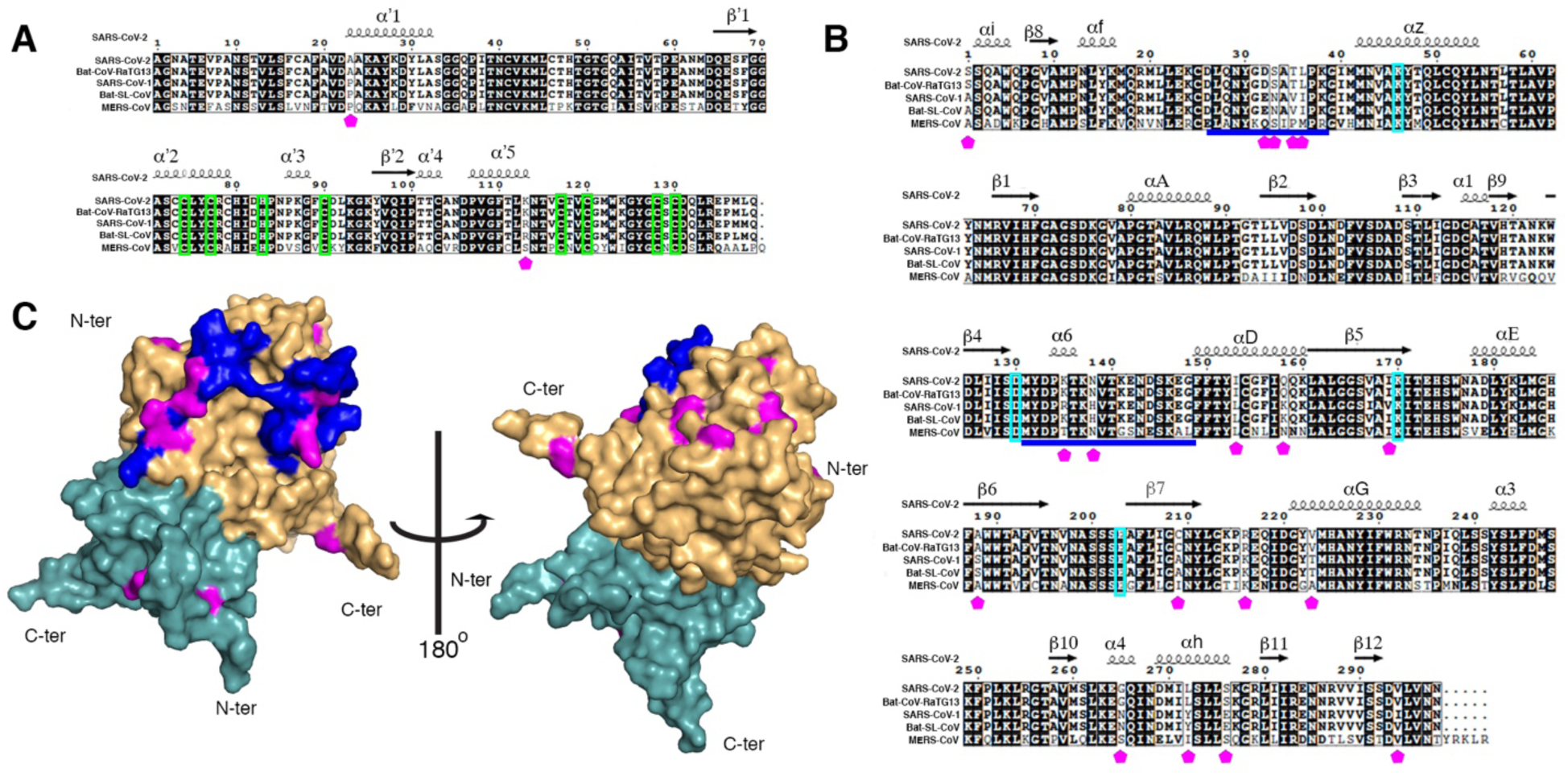
Sequence alignment of SARS-CoV-2 with other coronavirus and the mutations mapped in the structure. Multiple sequence alignment of nsp10 (A) and nsp16 (B). Black shading represents 100% residue identity and the punctual mutations are indicated with the magenta pentagon. In nsp10, the green boxes indicate the Zn binding site. In nsp16, the flexible loops involved in Cap and SAM binding stability and differ from SARS-CoV-1, and/or MERS-CoV are underlined in blue and the cyan boxes indicate the catalytic KDKE. C, Cartoon representation of the structure of SARS-CoV-2, with nsp10 subunit in teal and nsp16 in tan. The single amino acid changes between SARS-CoV-1 and SARS-CoV-2, and the residues comprising nsp16 flexible loops are mapped on the surface projection, as in (A) and (B).

Similarly, nsp16 from SARS-CoV-2 is 100% identical to Bat-CoV-RaTG13, 93% identical to Bat-SL-CoV Rs4247, and 95% identical to SARS-CoV-1 (Figure 3B), but only 66% identical to MERS-CoV. SARS-CoV-2 nsp16 contains the highly conserved K46, D130, K170, E203 (KDKE) motif for methyl-transfer found in many 2’-O-MTases (22), including the more phylogenetically distant MERS-CoV. Furthermore, we confirmed that residues N43, Y47, G71, A72, S74, G81, D99, N101, L100, D114, and M131, which coordinate SAM substrate binding in SARS-CoV-1 through hydrogen bonds and water-mediated interactions (4), are also 100% conserved in SARS-CoV-2 (Fig. 2F). The high conservation of nsp10 and nsp16 sequences, and the identification of 100% conserved catalytic and substrate-binding residues suggest that SARS-CoV-2 nsp10-nsp16 function and structure are also highly conserved.

### Structural comparison of 2’-O-MTase from SARS-CoV-2 with SARS-CoV-1 and MERS-CoV

In the primary sequence alignments of nsp10, there are only two amino acid differences (A34P and K113) between SARS-CoV-2 and SARS-CoV-1. These changes do not affect the structure of nsp10 or its interactions with Zn^2+^ or nsp16. On the other hand, there are 18 amino acid differences between the SARS-CoV-2 and SARS-CoV-1 nsp16 sequences (Figure 3 A and B). Many of these amino acid variations in SARS-Cov-2 occurred on the solvent-exposed surface (Figure 3C), and are localized in the most flexible loops that surrounds the nucleotide binding site comprised of amino acids 26-38. This difference could have steric implications in the RNA binding, especially for the first loop where four mutations are located. The amino acid variant of Glu-32 to Asp and Ile-37 to Leu are homonymous mutations; however, Asn-33 to Ser and Val-35 toThr might have steric implications, since these two residues together with Leu-37 are nearby the nsp10-nsp16 interface. The other mutations are located in the N-terminal and the β-sheet. However, structural and catalytic studies are required in order to understand the role of these mutations.

Structural alignment using the DALI server shows the highest coverage, percent identity, and structural conservation with the SARS-CoV-1 complex bound to SAM (PDB code 3r24 (4), z= 48.3, r.m.s.d=0.9) or to S-adenosylhomocysteine (PDB code 2xyq (4) z=47.5, r.m.s.d.=1). The structure is also highly similar to the MERS-CoV complex in the apo form (PDB code 5yn5 (29), z=47.2, r.m.s.d.=0.8) and with SAM bound (PDB code 5YN6 (29), z=46.7, r.m.s.d.= 0.7). The differences found between the structures is mostly influenced by the absence of some residues from the N and C-terminal sequences and the flexibility of the cap-binding loop at residues 26-38 and the loop 131-148 that forms part of both the cap binding groove and SAM cleft (Figure 3C), both of which are disordered in the MERS-CoV nsp16 structures (PDB codes 5yn5 and 5yn6 (29)) or have a different conformation in the structure of SARS-CoV-1 bound to SAM (PDB code 3r24).

Overall, the structure of SARS-CoV-2 nsp10-nsp16 heterodimer is very conserved among coronaviruses and further supports the idea that there is functional and mechanistic conservation of the nsp16 2’-O-MTases for mRNA maturation in coronaviruses.

### Comparison with other eukaryotic MTases

Although the primary sequences vary, MTases of viral MTases have high similarity with eukaryotic MTases (28,30). To identify structurally-related, non-viral MTases, the nsp16-nsp10 structure was submitted to the DALI server (30), which identified eukaryotic MTases that are structurally homologous. These include the human cap-specific 2’-O-MTase (PDB code 4n48, z=13.7, r.m.s.d= 3.4) and the yeast 27S pre-rRNA guanosine (2922)-2’-O)-MTase (PDB code 6em5-W, z=12.4, r.m.s.d.=3.1). The main differences are located in the loops that surround the beta sheet that is fully conserved, which corroborates the high degree of structural conservation in this pathway. The close alignment reveals that the mechanism of coronavirus 2’-O-methylation is conserved with eukaryotic cell mechanisms.

## Conclusions

The SARS-CoV-2 pandemic has yielded a world-wide effort to understand the molecular mechanisms involved in virus transmission, virulence, and replication. The ultimate goal is to identify viral proteins that are amenable for drug targeting and epitopes suitable for vaccine development. Although, previous studies conducted in SARS-CoV-1 and MERS-CoV enlighten the pathway for drug discovery and vaccine development, no approved treatments were fully developed. In addition, due to the high replication error rate of ssRNA viruses, small amino acid variations could cause resistance to previously described drugs or antibodies, as well as increase the pathogenicity. Thus, in order to ensure an accurate approach for drug discovery, the CSGID implemented a structural genomics pipeline to solve the structure of proteins essential for the replication of SARS-CoV-2. Here we presented the structure of the 2’-O-MTase, nsp10-nsp16 heterodimer in complex with SAM at 1.8 Å resolution, which shares 98% identity with the 2’-O-MTase of SARS-CoV-1. All the amino acid variations in the new SARS-CoV-2 were located at the solvent exposed surface of the heterodimer, and not among the residues of the catalytic site, substrate binding site, or interface between nsp10-nsp16. Furthermore, the only identified discrepancies in our model versus previous models were in the flexible loop regions 28-38 and 131-148 and the N- and C-terminal residues. As the 2’-O-MTase activity of nsp10-nsp16 is required for viral replication-transcription in host cells, these structural data can be used for developing new specific treatments against SARS-CoV-2. Importantly, the structure reported here is the highest resolution structure thus far of a coronavirus 2’-O-MTase and is amenable for computational modeling of potential inhibitors.

## Materials and Methods

In order to increase the probability to obtain high quality crystals suitable for structure determination, the CSGID implemented two different approaches, which are listed here and identified by their PDB accession code.

### Method 1, PDB code 6wh4

#### Cloning and protein production

The genes from nsp10 and nsp16 were codon optimized for expression in *E. coli*, synthetized (Twist Bioscience) and cloned into pMCSG53 (31) vector, which contains a TEV cleavable N-terminal 6xHis-tag, encodes ampicillin resistance, and genes for rare codons. The plasmids were transformed into *E. coli* BL21(DE3)(Magic) cells (32). The proteins were expressed separately starting with 10 ml LB supplemented with 130 μg/ml and 50 μg/ml KAN overnight culture at 37°C and 220 rpm. The next day, 3 liters of TB medium supplemented with 200 μg/ml AMP and 50 μg/ml KAN were inoculated at 1:100 dilution with the overnight culture and incubated at 37°C and 220 rpm. Protein expression was induced at OD_600_=1.8-2 by addition of 0.5 mM isopropyl β-d-1-thiogalactopyranoside and the cultures were further incubated at 25°C, at 200 rpm for 14 h (33). The cells were harvested by centrifugation and resuspended (1g of cells: 5 ml of lysis buffer) in lysis buffer (50 mM Tris, 0.5 M NaCl, 10% glycerol, 0.1% IGEPAL CA-630) and frozen at −30°C until purification.

#### Protein purification

Frozen pellets were thawed and sonicated at 50% amplitude, in 5 s x 10 s cycle for 20 min at 4°C. The lysate was cleared by centrifugation at 18,000 x *g* for 40 min at 4°C, the supernatant was collected, and the protein was purified as previously described with some modifications (34). The supernatant was loaded into a His-Trap FF (Ni-NTA) column using a GE Healthcare ÅKTA Pure system in loading buffer (10 mM Tris-HCl pH 8.3, 500 mM NaCl, 1 mM Tris(2-carboxyethyl) phosphine (TCEP), 2 mM MgCl_2_ and 5% glycerol). The column was washed with 10 column volumes (cv) of loading buffer and 10 (cv) of washing buffer (10 mM Tris-HCl pH 8.3, 500 mM NaCl, 25 mM imidazole), and was eluted with elution buffer (10 mM Tris pH 8.3, 500 mM NaCl, 1 M imidazole). The protein was loaded onto a Superdex 200 26/600 column and ran with loading buffer. The protein was collected and mixed with TEV protease (1:20 protein:protease) and incubated overnight at room temperature for removal of the 6xHis-tag. The cleaved tag was separated from protein by Ni-NTA-affinity chromatography using loading buffer. Nsp10 was collected in the flow through, whereas nsp16 eluted at 50 mM imidazole. To form the nsp10-nsp16 complex, the pure cleaved proteins were mixed at a 1:1 molar ratio at approximately 2 mg/ml in loading buffer and incubated for 1 h, then dialyzed in crystallization buffer (10 mM Tris-HCl pH 7.5, 150 mM NaCl 5% glycerol) for 2 h (23). SAM was added to a final concentration of 2 mM. The complex was concentrated to 5.5 mg/ml and set up for crystallization immediately.

#### Crystallization and data collection

The pure nsp10-nsp16 complex (1:1 ratio for nsp10 and nsp16) + SAM was set up as 2 µl crystallization drops in (1µl protein:1µl reservoir solution) in 96-well plates (Corning) and equilibrated using commercially available Classics II, PEG’s II, Anions and ComPas Suites (Qiagen). Diffraction quality crystals appeared after 5 days in condition A7 of the ComPAS screen (0.2 M Calcium acetate, 0.1 M HEPES, pH 7.5, 18% w/v PEG 8000).

The data set was collected on the LS-CAT 21-ID-F beamline at the Advanced Photon Source (APS) at the Argonne National Laboratory. A total of 200 images, which corresponded to 120 degrees of the spindle axis rotation were collected. Images were indexed, integrated and scaled using HKL-3000 (35).

#### Structure solution and refinement

The structure of native nsp10-nsp16 complex crystal from SARS-CoV-2 was solved by Molecular Replacement with Phaser (36) from the CCP4 Suite (37) using the crystal structure of the nsp10/nsp16 heterodimer from SARS (PDB ID 3r24) as a search model. Initial solutions went through several rounds of refinement in REFMAC v. 5.8.0258 (38) and manual model corrections using Coot (39). The water molecules were generated using ARP/wARP (40), S-Adenosylmethionine (SAM), Zinc ions and ligands were added to the model manually during visual inspection in Coot. Translation–Libration–Screw (TLS) groups were created by the TLSMD server (41) (http://skuldbmsc.washington.edu/~tlsmd/) and TLS corrections were applied during the final stages of refinement. MolProbity (42) (http://molprobity.biochem.duke.edu/) was used for monitoring the quality of the model during refinement and for the final validation of the structure. Final model and diffraction data were deposited to the Protein Data Bank (https://www.rcsb.org/) with the assigned PDB code 6wh4.

### Method 2 PDB code 6w61

#### Cloning and protein production

The cloning strategy for nsp10 and nsp16 was the same as the previous protocol except that the plasmids were transformed into BL21(DE3)-Gold strain cells. For protein expression, 4 L culture of LB Lennox medium was grown at 37°C (190 rpm) in presence of 150 μg/ml ampicillin to OD_600_ ∼1.0, then incubated at 4°C until the cultures reached 18°C.Protein production was induced by addition of 0.2 mM IPTG, 0.1% glucose, 40 mM K_2_HPO_4_ to the medium, and cultures were incubated at 18 °C for 20 hours. Bacterial cells were harvested by centrifugation at 7,000 x *g* and cell pellets were resuspended in lysis buffer (500 mM NaCl, 5% (v/v) glycerol, 50 mM HEPES pH 8.0, 20 mM imidazole and 10 mM β-mercaptoethanol).

#### Protein purification

Purification of nsp10 and nsp16 (PDB ID 6W61) from the fresh suspension of cells in lysis buffer was sonicated at 120W for 5 minutes (4s ON, 20 sec OFF cycle). The cellular debris was removed by centrifugation at 30,000 *g* for one hour at 4°C. The supernatant was mixed with 4 ml of Ni^2+^ Sepharose (GE Healthcare Life Sciences) equilibrated with lysis buffer supplemented to 50 mM imidazole pH 8.0 and suspension was applied on Flex-Column (420400-2510) connected to Vac-Man vacuum manifold (Promega). Unbound proteins were washed out via controlled suction with 160 ml of lysis buffer (50 mM imidazole). Bound proteins were eluted with 20 ml of lysis buffer supplemented to 500 mM imidazole pH 8.0. 2 mM DTT was added followed by TEV protease treatment at 1:20 protease:protein ratio at 4°C overnight. Next, proteins were concentrated and run on size exclusion column Superdex 200 in lysis buffer supplemented with 1 mM TCEP. The fractions containing protein of interest were pooled and run through Ni^2+^ Sepharose and the proteins were collected in the flow through. Finally, buffer was exchanged to crystallization buffer (150 mM NaCl, 20 mM HEPES pH 7.5, 1 mM TCEP) (23) via 10X concentration/dilution repeated 3 times. Final concentrations of Nsp10 and Nsp16 were 8.5 mg/ml and 5 mg/ml, respectively. Proteins were either flash cooled or used immediately for crystallization trials.

#### Crystallization and data collection

To obtain crystal form 2, the complex was screened by the sitting-drop vapor-diffusion method in 96-well CrystalQuick plates (Greiner Bio-One) which had been set up using Mosquito liquid dispenser (TTP LabTech), (400nl protein: 400 nl reservoir solution) equilibrated against 135 nl reservoir solution against MCSG1, MCSG4 (Anatrace), SaltRX (Hampton) and INDEX (Hampton) screens were used for protein crystallization at 16°C. Diffraction quality crystals appeared after 3 days in MCSG4 G4 (0.1 M Sodium citrate pH 5.6, 10 %(w/v) PEG4000).

Data were collected at the SBC ID beamline sector 19 at the APS. A total of 954 diffraction images from five different parts of the rod-shaped crystal were collected to 2.00 A on the Dectris Pilatus3 × 6M detector. The data were processed and scaled by HKL3000 (43).

#### Structure solution and refinement

The structure was determined by molecular replacement using MolRep (44) implemented in HKL3000 and 6w4h structure as a search model. The structure was refined iteratively using Phenix.refine (45) and manually corrected in Coot. Water molecules were automatically generated in Phenix.refine while SAM, Zinc ions and ligands were added manually in Coot. The stereochemistry of the structure was validated using MolProbity and the structure was deposited to PDB with the assigned PDB code 6w61.

#### Structural and sequence alignment

The protein sequence of nsp10 and nsp16 from SARS-CoV-2 (YP_009725295.1), Bat-CoV-RaTG13 (QHR63299.1) and Bat-SL-CoV Rs4247 (ATO98179.1) SARS-CoV-1 (ACZ72252.1) and MERS-CoV (YP_009047238.1) were obtained from the NCBI database. The multiple sequence alignment was performed using ClustalO (https://www.ebi.ac.uk/Tools/msa/clustalo/) and merged with the PDB file of 6w4h using ESPript 3.x (46). The PDB coordinates of the SARS-CoV-1 nsp10-nsp16 complex were analyzed on the DALI (47), POSA and (48) FATCAT (49) servers to perform structural and sequence alignment. Structural alignments and structure figures were conducted using Matchmaker in Pymol (50) open source V 2.1.

## Acknowledgements

This project has been funded in whole or in part with Federal funds from the National Institute of Allergy and Infectious Diseases under Contract Nos. HHSN272201200026C and HHSN272201700060C and Grant Nos. U01AI124316 and R01 GM05789 and from the National Institute of General Medical Sciences grant GM118187, both from National Institutes of Health, Department of Health and Human Services. This research used resources of the Advanced Photon Source, a U.S. Department of Energy (DOE) Office of Science User Facility operated for the DOE Office of Science by Argonne National Laboratory under Contract No. DE-AC02-06CH11357. Use of the LS-CAT Sector 21 was supported by the Michigan Economic Development Corporation and the Michigan Technology Tri-Corridor (Grant 085P1000817).

---

| PDB Accession Code | 6w4h | 6w61 |
| --- | --- | --- |
| **Crystallization Conditions** |  |  |
| Screen conditions | 0.2M Calcium acetate, 0.1M HEPES pH 7.5,<br>18% (w/v) PEG8000 | 0.1 M sodium citrate, pH 5.6,<br>10% (w/v) PEG4000, 10%(w/v) isopropanol |
| Protein concentration (mg/ml) | 5.3 | 16.7 |
| **Data Collection** |  |  |
| Space group | *P3<sub>1</sub>21* | *P3<sub>1</sub>21* |
| Unit cell parameters (Å; °) | *a* = 167.7, *b* = 167.7, *c* = 51.9;<br>*α* = *β* = 90.0, *γ* = 120.0 | *a* = 168.6, *b* = 168.6, *c* = 52.0;<br>*α* = *β* = 90.0, *γ* = 120.0 |
| Resolution range (Å) | 30.00 – 1.80 (1.83 - 1.80) | 50.00 - 2.00 (2.04 - 2.00) |
| No. of reflections | 77,886 (3,873) | 57,010 (2,781) |
| *R*<sub>merge</sub> (%) | 6.0 (76.5) | 13.9 (233.0) |
| Completeness (%) | 100.0 (100.0) | 100.0 (100.0) |
| *⟨I/σ(I)⟩* | 29.3 (2.6) | 37.1 (1.0) |
| Multiplicity | 7.3 (6.8) | 6.2 (6.3) |
| Wilson *B* factor | 25.4 | 49.0 |
| **Structure Determination** |  |  |
| MR initial model (PDB ID) | 3r24 | 6w4h |
| **Refinement** |  |  |
| Resolution range (Å) | 29.76 - 1.80 (1.85 - 1.80) | 42.33 - 2.00 (2.04 - 2.00) |
| Completeness (%) | 100.0 (100.0) | 99.7 (96.3) |
| No. of reflections | 73,752 (5,432) | 56,859 (2,522) |
| *R*<sub>work</sub>/*R*<sub>free</sub>, (%) | 14.9/16.3 (22.9/23.9) | 17.2/19.3 (30.4/32.3) |
| Protein chains/atoms | 2/3,201 | 2/3,390 |
| Ligand/Solvent atoms | 65/442 | 6/141 |
| Mean temperature factor (Å<sup>2</sup>) | 31.5 | 41.0 |
| **Coordinate Deviations** |  |  |
| R.m.s.d. bonds (Å) | 0.004 | 0.002 |
| R.m.s.d. angles (°) | 1.183 | 0.473 |
| **Ramachandran plot** |  |  |
| Favored (%) | 97.0 | 96.8 |
| Allowed (%) | 3.0 | 3.2 |
| Outside allowed (%) | 0.0 | 0.0 |

